# An EcR probe reveals mechanisms of the ecdysone-mediated switch from repression-to-activation on target genes in the larval wing disc

**DOI:** 10.1101/2022.04.07.487542

**Authors:** Joanna Wardwell-Ozgo, Douglas Terry, Colby Schweibenz, Michael Tu, Ola Solimon, David Schofeld, Kenneth Moberg

## Abstract

Fluctuating levels of steroid hormones provide both systemic and local cues to synchronize metazoan development and control germline and homeostatic processes. The main steroid hormone in *Drosophila* is ecdysone (Ec), which upon binding of its active form (20E) converts its receptor, EcR, from a transcriptional repressor to activator. Multiple co-repressors and co-activators are proposed to act with EcR in different tissues to control diverse targets and processes, including apoptosis, cell migration, and proliferation. Despite these diverse roles, relatively little is known regarding how EcR translates Ec temporal gradients into modulation of individual target genes. Here we use an Ec-binding fragment of EcR (EcR^LBD^) as a ‘sponge’ to sequester coregulators and probe the state of EcR activity as larval wing cells traverse the 3^rd^ instar Ec gradient. This approach reveals a dramatic and rapid shift from EcR mediated repression-to-activation in late L3 cells, and that the extent of repression varies between targets. An Ala^483^Thr mutation that disrupts binding of the co-repressor Smr compromises the ability of EcR^LBD^ to derepress reporters, but also limits its ability to block activation, suggesting either that a coactivator shares an EcR-interaction interface with Smr or that Smr-repression primes targets for 20E activation. Molecular and genetic data reveal that EcR^LBD^ sequesters 20E, and that EcR^LBD^ phenotypes can be modulated by manipulating intracellular 20E levels with Ec importer (EcI) and Cyp18a1, which inactivates 20E. Finally, we provide evidence that Smr repression of EcR activity varies spatially and by target in the wing disc. In sum these data reveal that relief of EcR-Smr repression is a major contributor to 20E induction of EcR targets in larval wing discs and highlight EcR^LBD^ as an effective probe to define EcR-20E gene regulatory mechanisms *in vivo*.

## Introduction

The growth and patterning of developing metazoan tissues is coordinated by signals that include adhesion events between neighboring cells, secreted morphogens and ligands that act regionally within a tissue or compartment, and systemic inputs from nutrients and hormones that act at the level of the whole organism. These latter hormonal inputs synchronize growth rates and coordinate timing of developmental events across multiple tissues, and can take the form of peptides (e.g., in the case of insulin-like peptides) or cholesterol-derived steroids, which are well-known for triggering progression through major developmental transitions e.g., puberty [reviewed in 1, 2]. In the fruit fly *Drosophila melanogaster*, the main steroid hormone ecdysone (Ec) is synthesized in the larval prothoracic gland (PG) and secreted into the hemolymph, which theoretically bathes all internal tissues in an equivalent Ec concentration [reviewed in 1]. Ec can be imported into cells by members of the Oatp-class of organic ion transporters and then converted intracellularly into its bioactive form, 20-hydroxyecydsone (20E) [3-5].

The 20E ligand elicits changes in gene expression by interacting with the ligand-binding domain (LBD) of the ecdysone receptor (EcR), a type-II nuclear hormone receptor (NHR) that binds a defined palindromic DNA sequence as a heterodimer with its partner an RxR homolog Ultraspriacle (Usp) [6-10]. EcR-Usp complexes can repress targets in the absence of 20E and activate targets as 20E titers rise [11-14]. Like other NHRs, this EcR functional plasticity is presumably based on 20E-induced allosteric changes in the LBD which evict corepressors and allow binding of coactivators [15]. Dynamic switches from repression to activation allow EcR to translate rising 20E concentrations into tissue- and stage-specific transcriptional programs that vary based on EcR isoforms present in each cell type, their patterns of genomic occupancy, and the spectrum of co-expressed EcR activators and repressors [16-21]. Through process, EcR contributes to diverse developmental processes, including apoptosis of polyploid larval cells [22, 23], proliferation of larval histoblast nests [24, 25], collective migration of cells in the ovary [26], and transitions between developmental stages, e.g., embryo-to-larva and larva-to-pupae [rev. in 27].

The temporal patterns of EcR transcriptional activity in developing tissues are predicted to mirror a series of developmentally programmed Ec peaks that begin in the embryo and conclude in the pupal phase [28]. These temporal gradients of Ec also have the potential to drive different expression patterns for low and high-threshold 20E-EcR targets in a manner akin to classic spatial morphogen gradients (e.g., Dpp [29]). Consistent with this hypothesis, recent studies show that EcR dynamically redistributes across the genome of late third instar wing imaginal disc cells as 20E titers rise, implying that EcR targets shift as development proceeds, even within a single cell type [30]. One potential interpretation of this data is that enhancers occupied by EcR only at low Ec titers are primarily regulated by EcR repression, while those only occupied at high Ec titers are primarily regulated by EcR activation. In addition, loci differentially bound by EcR across the timescale of these experiments had a lower motif density and EcR binding strength [30], suggesting that EcR recruitment to these temporally dynamic promoters might rely on cofactor recruitment or interactions with other transcription factors bound at shared enhancers. Testing these models of EcR mediated activation and repression on specific target genes, especially in rapidly dividing imaginal epithelial cells, is challenging given that *EcR* loss-of-function alleles or inhibition of Ec production in the PG each lead to developmental arrest [31, 32]. A more targeted approach of EcR RNAi depletion from its bound genomic sites is predicted to relieve EcR-mediated repression but also block activation, making it difficult to extrapolate roles of EcR-coregulator complexes on their targets. Similarly, overexpressed EcR variants (e.g., the *UAS-EcR*^*F645A*^ dominant negative [14]) are predicted to promiscuously bind EcR target sites throughout the genome and overwhelm specificity mechanisms.

To enable targeted analysis of the role of endogenous EcR and 20E-dependent co-regulators in controlling specific target genes in intact tissues, we have created a *UAS*-transgene encoding a fragment of the *EcR* coding sequence encompassing the LBD domain (EcR^LBD^). This EcR^LBD^ fragment is predicted to bind 20E and co-regulators that interact with the activation function-2 domain (AF2), which in embedded in the LBD, and engage in 20E-regulated interactions with coregulators. This approach is based on a prior study suggesting that this fragment could act as a dominant negative EcR *in vivo* [33]. The *UAS-EcR*^*LBD*^ system enables spatiotemporally controlled disruption of LBD-dependent interactions between endogenous EcR and coregulators, and visualization of downstream effects on expression of candidate targets. When expressed in L3 wing discs, EcR^LBD^ reveals a temporal switch from EcR-mediated repression to activation that occurs in a narrow temporal window within the L3 stage. Significantly, EcR^LBD^ induction of heterologous EcR reporters in young L3 wing discs is genetically insensitive to depletion of the *Oatp74D* ecdysone importer EcI, indicating that EcR^LBD^ titrates a corepressor. EcR^LBD^ repression of these same reporters in late L3 wing cells is blocked by depletion of *Cyp18a1*, which inactivates 20E [34, 35], indicating that EcR^LBD^ repression can be overcome by elevating available 20E. Consistent with these effects, we find that EcR^LBD^ binds 20E in cells, indicating that it likely retains allosteric mechanisms that exchange repressors and activators following 20E binding. Finally, we use the EcR^LBD^ fragment as a platform to test the effect of the Ala^483^Thr mutation (A483T), which blocks interaction with the Smrter (Smr) corepressor [36]. Relative to wildtype EcR^LBD^, A483T is defective in induction of EcR targets in young L3 wing cells, including the Broad transcription factor, and, unexpectedly, is also defective in repression of endogenous EcR activity in late L3 wing cells, suggesting that 20E-induced eviction of Smr may result in binding of coactivators to EcR using an interaction interface shared by Smr. In sum, these studies reveal a temporal switch between EcR repression and activation in L3 larval wing cells that is sensitive to genetic manipulation of local 20E levels and connect these mechanisms to EcR-Smr dependent repression of physiologic EcR targets, such as Broad. These data validate the EcR^LBD^ fragment as an effective tool to assess the relative contribution of EcR repressors and activators to direct the temporal control of individual EcR targets in wing disc cells, and also provide proof-of-principle data with A483T showing that effects of *EcR* alleles can be effectively modelled in the EcR^LBD^ system.

## Results

### *EcR*^*LBD*^ structure and effects on 20E-regulated developmental processes

To enable targeted disruption of interactions between EcR and co-factors in specific cells, a fragment of full-length *EcR* cDNA encoding the hinge, ligand binding domain, and C-terminal end [regions D, E, F according to nomenclature in [14]] was tagged with a C-terminal red fluorescent protein tag (nls-RFP) and inserted into a *UAS* transgene (*UAS-EcR*^*LBD*^) (**Fig. 1A**). EcR^LBD^ contains the 20E binding region and the Activation Function-2 domain (AF2), which seeds interactions with activators and repressors and contains the presumed dimerization interface with the EcR partner and RXR homolog Ultraspiracle (Usp) [37]. EcR^LBD^ lacks the DNA binding domain and the isoform-specific A/B regions, which include the transactivation domain Activation Function-1 (AF1). Several mutations that affect EcR transcriptional activity lie within the EcR^LBD^ region (annotated in **Fig. 1A**), including *Ala*^*483*^*Thr* (A483T), which lies within the ‘Ti’ domain and disrupts interaction with the fly NCoR1 repressor homolog Smrter (Smr) [36] and *Phe*^*645*^*Ala* (F645A), which is within AF2 and blocks 20E-stimulated gene expression [14, 38]. Significantly, ubiquitous expression of a similar EcR-LBD fragment using a heat shock-inducible promoter elicits dominant-negative effects *in vivo* [33], likely by sponging EcR regulators.

**Figure 1.**
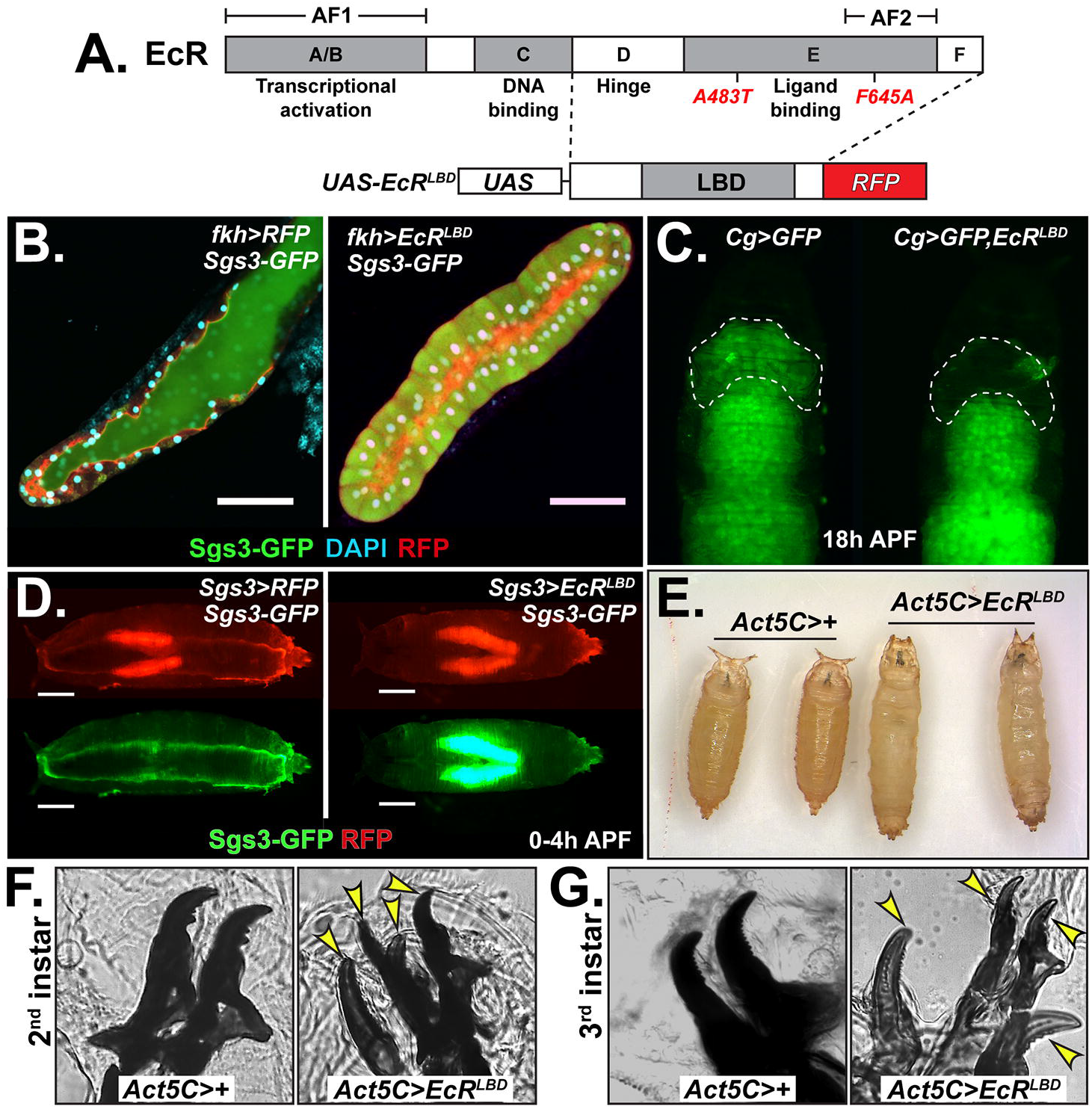
EcR^LBD^ disrupts 20E-EcR regulated processes. (**A**) The EcR^LBD^ consists of the hinge, ligand binding domain (LBD) and F regions with a C-terminal RFP fusion. AF1 and AF2 domains are indicated. Mutations that disrupt EcR functions are indicated (red). (**B**) Relative to control (*fkh>RFP*), EcR^LBD^ (*fkh>EcR*^*LBDi*^;red) prevents luminal secretion but not production of Sgs3 glue protein (*Sgs3-GFP*;green) in salivary gland cells. (**C**) Relative to control (*Cg>GFP*), EcR^LBD^ (*Cg>EcR*^*LBD*^,*GFP)* has minimal effect on FB remodeling at 18h APF but blocks FB migration into the pupal head (dashed outlines). (**D**) Relative to control (*Sgs3>RFP*), EcR^LBD^ (*Sgs3>EcR*^*LBD*^) blocks Sgs3 glue (*Sgs3-GFP*) expectoration at 0-4h APF. Ubiquitous EcR^LBD^ causes (**E**) elongated pupae, and mouth hook retention in (**F**) 2nd and (**G**) 3rd instar mouth hooks.

To test the prediction that *UAS-EcR*^*LBD*^ sponges key EcR cofactors and perturbs 20E-regulated processes, the effect of EcR^LBD^ expression was tested on a group of EcR-regulated processes: glue protein production, secretion, and expectoration in the larval and pupal salivary glands (SGs), fat body (FB) migration into the pupal head case, and mouth hook molting between the larval instars (**Fig. 1B-G**). Expression of *EcR*^*LBD*^ in SGs with *fkh-Gal4* blocks EcR-regulated secretion of a GFP-tagged Sgs3 glue protein (*Sgs3-GFP*) [39-41] into the lumen of larval SGs but does not block Sgs3-GFP production (**Fig. 1B**). By comparison, *fkh-Gal4* driven RNAi knockdown of *EcR*, which will remove EcR from sites of genomic occupancy, or RNAi of the 20E-importer *EcI* [42], which will selectively block 20E-induced gene expression, also block lumenal secretion in larval SGs, with cell-to-cell variability in blocking Sgs3-GFP production (**Fig. S1C**). *fkh-Gal4* transgenic expression of *UAS-EcR*^*F645A*^, a dominant negative protein predicted to fill EcR binding sites across the genome, also blocks Sgs3-GFP lumenal secretion in larval SGs, with some cells also failing to express Sgs3-GFP (**Fig. S1B**). Expression of *EcR*^*LBD*^ in pupal SGs (0-4hr after puparium formation; APF), using the *Sgs3-Gal4* driver [14] completely blocks Sgs3-GFP in a manner similar to *EcR*^*RNAi*^, *EcR*^*F645A*^, or overexpression of the *Cyp18a1* P450 enzyme that inactivates 20E [34, 35] (**Figs. 1D** and **S1D-E**). Together, these data indicate that *EcR*^*LBD*^ specifically disrupts some EcR-dependent processes in larval SG cells but not others. These different phenotypic consequences of EcR^LBD^ versus EcR or EcI loss, or EcR^F645A^ overexpression, may discriminate developmental events controlled by LBD and AF2 interactions with activators and repressors, versus those that respond to loss of EcR occupancy at genomic targets, a deficit in 20E-driven activation, or forced binding of EcR^F645A^ to sites throughout the genome.

In response to 20E-EcR signaling, larval FB sheets disassemble into individual cells that relocalize throughout the animal, including into the pupal head case [14, 43]. To assess effects of EcR^LBD^ on FB remodeling and localization, the *Cg-Gal4* driver [44] was used to drive the *UAS-EcR*^*LBD*^ transgene in FB cells. *Cg-Gal4* expression of *EcR*^*LBD*^prevents migration of FB cells into the pupal head but does not prevent disassembly of FB sheets, as indicated by the evenly packed FB cells visible in *Cg>GFP* controls and *Cg>EcR*^*LBD*^ pupae (**Fig. 1C**). By contrast, *Cg-Gal4* driven expression of either *EcR*^*F645A*^, *EcI*^*RNAi*^, or *EcR*^*RNAi*^ in the larval FB inhibits both migration and remodeling, as indicated by the paired FB sheets visible in whole larvae **(Fig. S1F**). Ubiquitous expressing *EcR*^*LBD*^ from the *Act5c-Gal4* driver prevents pupation of most animals and is fully lethal (**Fig. S1G**); however, survivors pupate on a delayed timeline and fail to shorten the head-to-tail axis relative to control pupae (**Fig. 1E**), which resembles phenotypes of *EcR* loss-of-function alleles [reviewed in 1]. Among *Act5c>EcR*^*LBD*^ larvae, ubiquitous EcR^LBD^ expression is associated with retention of 1^st^ instar mouth hooks in 5% of 2^nd^ instar larvae (n=22). However, 53% of 3^rd^ instar larvae retain 2^nd^ instar mouth hooks and of these, 27% have extra mouth hooks and 26% also retain 2^nd^ instar molted cuticle (n=22) (arrows in **Fig. 1F,G**). Together, these FB and mouth-hook data provide further evidence that EcR^LBD^ can disrupt 20E-EcR signaling in multiple tissues and at various developmental stages, suggesting that this fragment may sequester critical EcR interactors, including co-regulators that interact with the AF2 domain.

### EcR^LBD^ reveals a repression-to-activation switch in larval wing discs

During the L3 larval stage, systemic 20E levels fluctuate in three small peaks that are followed by a large peak that triggers the pupal transition [reviewed in 45]. At a genomic level, passage through the larval-to-pupal transition is associated with a shift in sites of genomic binding by EcR that in turn correlate with shifts in patterns of repression or activation of EcR target genes [14, 30, 46]. However, whether individual targets are predominantly repressed or activated by EcR, and the identity of EcR-cofactors involved in these mechanisms, remain undefined.

As a first test of the effect of EcR^LBD^ on individual target genes *in vivo*, two EcR reporter lines containing EcR response elements (EcRE) extracted from their endogenous loci, *EcRE-lacZ* (from the *hsp27* promoter, [47]) and *7xEcRE-GFP* (from the *EcR* promoter, [6]), were used to track endogenous EcR activity in developing wing discs as they traverse the temporal 20E gradient from early 3^rd^ larval instar (L3) to late-L3 just before entry in white prepupal (WPP) stage (**Fig 2A-C, G-I** and **S2A-B**). In otherwise wild type discs, *EcRE-lacZ* and *7xEcRE-GFP* are expressed at comparatively low levels throughout early-L3 wing discs (non-wandering), with regions of slightly higher expression either side of the dorsoventral (DV) boundary within the pouch. As these discs age through mid-L3 (early wandering) and late-L3 (late wandering/pre-WPP), both reporters become brighter and maintain higher expression in these pouch DV cells, especially in the *EcRE-lacZ* background. Relative to these control patterns (*en>RFP*), EcR^LBD^ elevates *EcRE-lacZ* and *7xEcRE-GFP* expression in early-L3 wing discs **(Fig. 2A** vs. **D** and **2G** vs. **J**) and represses these same reporters in mid- and late-L3 discs (**Fig. 2E-F** vs. **B-C** and **2K-L** vs. **H-I**). Significantly, the EcR^LBD^ repression in mid/late-L3 discs not only prevents the gradual rise in reporter activity normally seen at these stages (see **Fig. 2A-C**), but also represses both reporters down to near undetectable levels that are lower than normally seen in early-L3 discs (e.g., **Fig. 2E,K**). Analysis of timed discs reveals that this switch from EcR^LBD^-induced activation-to-repression of *EcRE-lacZ* has begun by approximately 146h after egg laying AEL (**Fig. 3A-C**). Heat-shock FLP-out wing clones expressing EcR^LBD^ in late-L3 discs also suppressed *EcRE-lacZ* activity (**Fig. S2C-D**). A majority of EcR^LBD^ protein within these clones localizes to the cytoplasm but a fraction overlaps the nuclear marker DAPI (**Fig. S2E-H**). Overall, these data indicate that *EcRE-lacZ* and *7xEcRE-GFP* undergo a switch from repressive inputs in mid-L3 wing discs to activating inputs in late-L3 wing discs, and that both mechanisms can be disrupted by EcR^LBD^.

**Figure 2.**
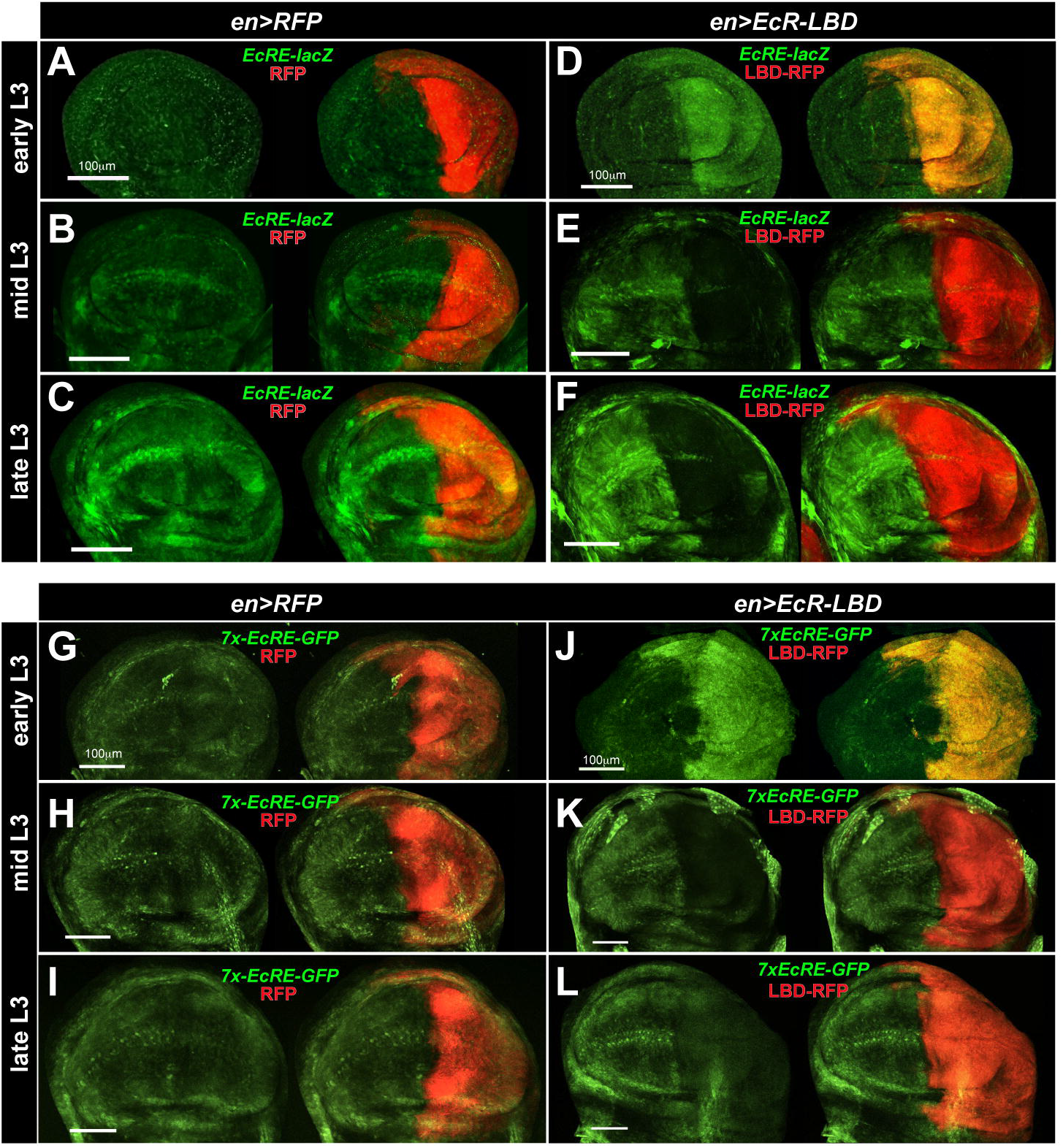
EcR^LBD^ reveals an EcR repression-to-activation switch in L3 wing discs. Optical (z-stack) projections of 3rd instar larval wing discs immunostained with anti-β- galactosidase (LacZ) to detect *EcRE-lacZ* expression (**A-F**) or anti-GFP to detect *7xEcRE-GFP* expression (**G-L**) in control (*en>RFP*; **A-C, G-I**) or *en>EcR-LDB* (**D-F, J-L**) discs. Note the progressive strengthening of both EcR activity reporters as discs age from early to late 3^rd^ instar and the sharp shift from EcR-LBD derepression to activation of both EcR reporters in the *en* domain (RFP-positive posterior half in all images) between early- and mid-3^rd^ instar discs (**D vs. E** and **J vs. K**).

**Figure 3.**
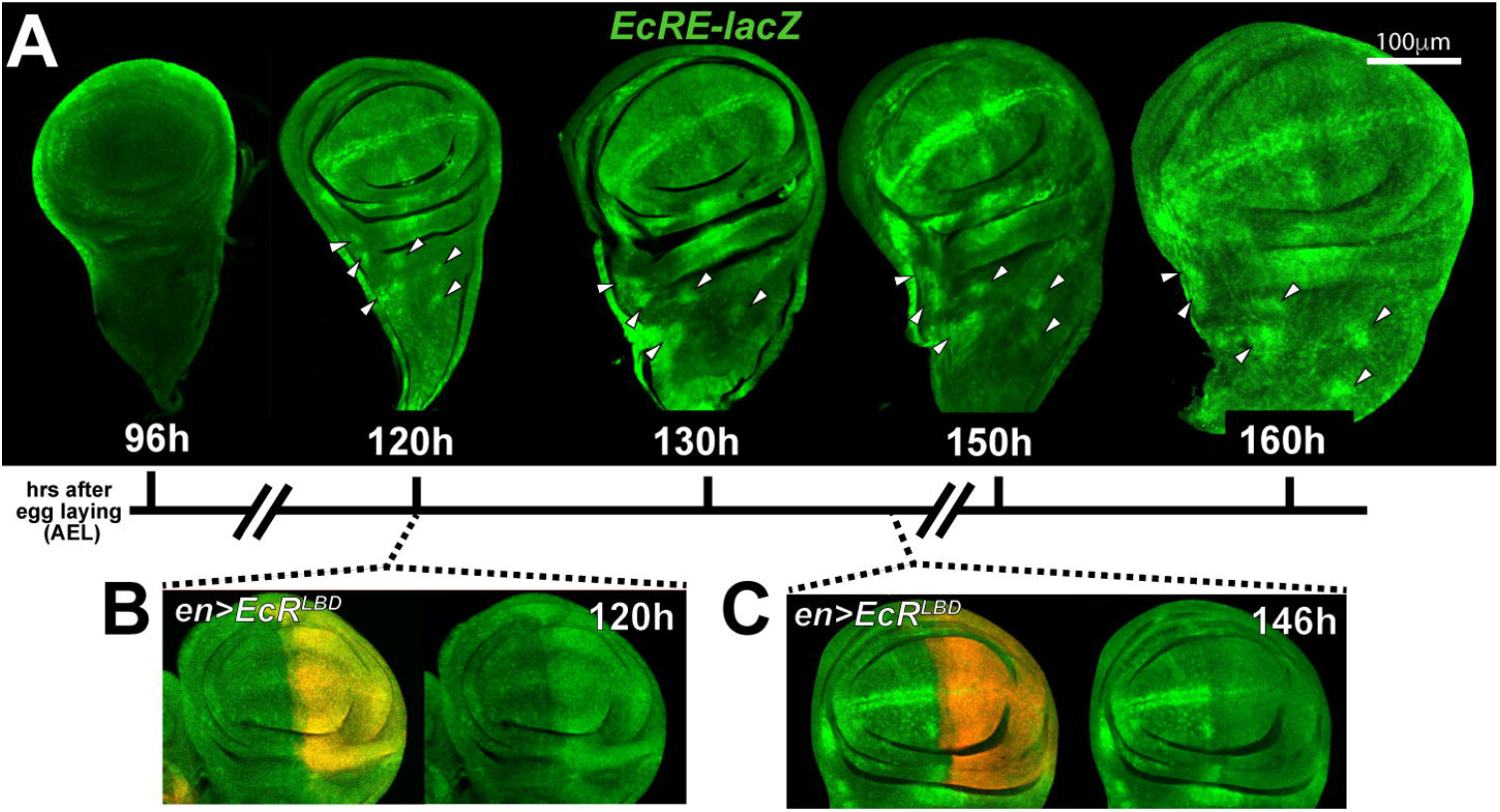
Timing the EcR repression-to-activation switch. (**A**) Optical projections of 3rd instar *EcRE-lacZ* larval wing discs immunostained with anti-β-galactosidase (green) at the indicated hours after egg laying (AEL). White arrows indicate presumptive SOP clusters. (**B**) At 120h AEL, *EcRE-lacZ* is activated by EcR^LBD^ (red), but (**C**) EcR^LBD^ has begun to repress *EcRE-lacZ* by 146h AEL.

### EcR^LBD^-mediated derepression and activation are genetically separable

The timing of the repression-to-activation switch on the *EcRE-lacZ* and *7xEcRE-GFP* reporters correlates with cessation of larval feeding and entry into the L3 wandering stage, which also parallels a rise in 20E levels that eventually peak in early pupae [27, 28]. To test the relationship between the repression-to-activation switch and feeding behavior, and to more precisely map its developmental timing, *EcRE-lacZ* expression patterns were analyzed at three different time points using the blue-dye food method [48]: larvae with ‘dark’ (i.e. full) guts are 12-24 hours before pupariation (L3^dark^=136-148h AEL); wandering larvae with ‘light’ guts have ceased feeding and are 6-12 hours before pupariation (L3^light^ =148-154h AEL); and wandering larvae with ‘clear’ (i.e. empty) guts are 1-6 hours before pupariation (L3^clear^ =154-159h AEL) (cartoon in **Fig. 4A**). When tested using this paradigm, *EcRE-lacZ* is elevated by expression of EcR^LBD^ (*en>EcR*^*LBD*^) in L3^dark^ wing cells but strongly repressed in L3^clear^ wing cells (**Fig. 4D,E** vs. **H**,**I**), suggesting that the repression-to-activation switch is complete by approximately 154h AEL and is closely timed with cessation of larval feeding. The comparatively low levels of 20E present in mid-L3 animals [49] suggest that EcR^LBD^-induced activation of *EcRE-lacZ* and *7xEcRE-GFP* at this stage may occur by sequestering of an EcR repressor rather than binding of 20E, while the reciprocal inhibitory effect of EcR^LBD^ in late-L3 disc discs is due to ‘sponging’ away 20E and/or a transcriptional activator. In support of this hypothesis, analysis of anti-RFP immunoprecipitates from lysates of *Actin>EcR*^*LBD*^ animals detects EcR^LBD^-associated 20E at 5-6 pg/mg (**Fig. 4B**), which approximates previous measurements of 20E in total L3 larval lysates [49]. To test the role of 20E in EcR^LBD^ transcriptional effects, intracellular 20E was manipulated with alleles of *EcI*, which imports 20E/Ec into cells [5], and *Cyp18a1*, which inactivates 20E by hydroxylation to 20,26-dihydroxyecdysone (see cartoon, **Fig. 4C**) [34, 35]. In mid-L3 wing discs, EcR^LBD^ induction of *EcRE-lacZ* is not blocked by *EcI* knockdown (*EcI*^*RNAi*^) or *Cyp18a1* overexpression (*Cyp18a1*^*oe*^) (**Fig. 4E** vs. **F-G**). As *en>Cyp18a1* is lethal (data not shown), this latter experiment used the *patched-Gal4* (*ptc-Gal4*) driver. These data suggest EcR^LBD^ activation of *EcRE-lacZ* in mid-L3 wing cells occurs mainly by loss of repression rather than 20E-triggered activation. Consistent with this hypothesis, *ptc-Gal4* expression of *Cyp18a1*^*RNAi*^ and *EcR*^*LBD*^ synergize to further elevate *EcRE-lacZ* among cells along the anterior:posterior (A/P) wing boundary (**Fig. S3A-C**). If EcR^LBD^ repression of *EcRE-lacZ* in late-L3 wing cells is due to sequestering ligand-dependent interactors (e.g., p160/Taiman, [26]), then supplementing intracellular 20E should restore *EcRE-lacZ* levels. To test this hypothesis, EcR^LBD^ was co-expressed with *EcI*^*oe*^ or *Cyp18a1*^*RNAi*^ in L3^clear^ discs (**Fig. 4H-K**). Compared to *en>EcR*^*LBD*^ alone, co-expression with *EcI*^*oe*^ or *Cyp18a1*^*RNAi*^ increased *EcRE-lacZ* activity to levels similar to *en>LBD* controls (yellow arrowheads in **Figs. 4H-I** vs. **J-K**), indicating that inhibition of EcR activity by *en>EcR*^*LBD*^ can be compensated elevated 20E. Interestingly, both *EcI*^*oe*^ and *Cyp18a1*^*RNAi*^ elevated *EcRE-lacZ* in wing pouch cells, but only *Cyp18a1*^*RNAi*^ was able to restore activity in bristle cells that flank the DV margin (asterisk, **Fig. 4K**), suggesting that 20E decay is more rate-limiting than Ec import in L3 wing cells. A similar rescuing effect on *EcRE-lacZ* activity is observed by co-expressing *Cyp18a1*^*RNAi*^ with *EcR*^*LBD*^ in the *ptc-Gal4* wing domain of late L3 wing cells (see arrows, **Fig. S3D-F**). Considered together, these epistatic interactions between *EcR*^*LBD*^ and *EcI* or *Cyp18a1* alleles suggest that *EcRE-lacZ* and *7xEcRE-GFP* are actively repressed in mid-L3 wing cells, and that this switches to 20E-dependent activation in wandering-stage L3 and late-L3 wing cells.

**Figure 4.**
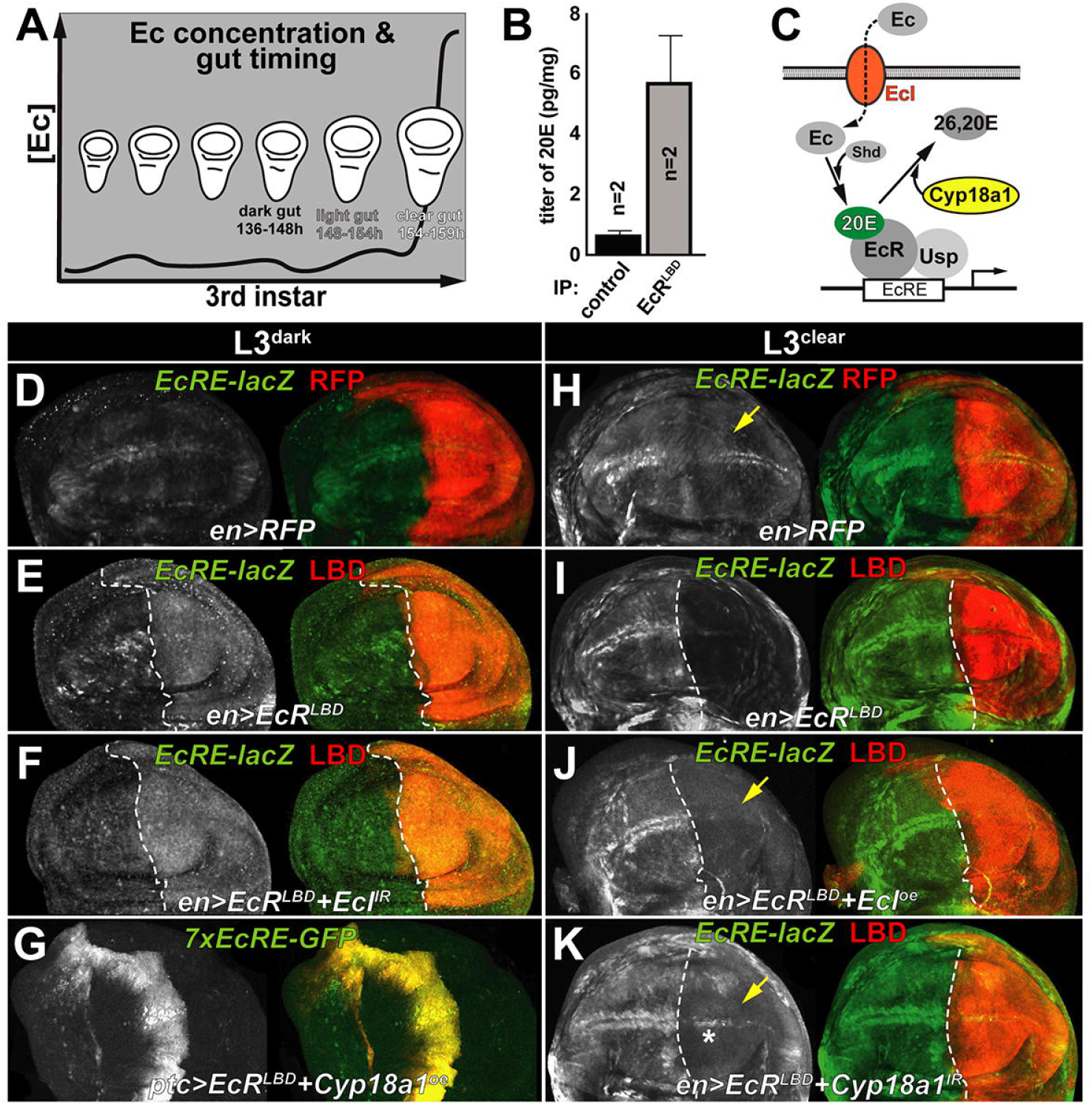
EcR^LBD^ derepression and activation can be distinguished by genetic dependence on EcI and Cyp18a1. (**A**) Cartoon schematic of 20E levels across the 3^rd^ instar and approximate timing of *dark, light* and *clear* guts. (**B**) Quantification of 20E hormone bound to anti-RFP immunoprecipitates from control (*actin>RFP*) or EcR^LBD^ expressing (*actin>EcR*^*LBD*^) L3 larvae. (**C**) Cartoon pathway of ecdysone (Ec) uptake into cells by EcI/Oatp74D importer (orange), conversion to 20E (green) by the cytochrome P450 enzyme Shade (Shd), and hydroxylation and inactivation by the Cyp18a1 enzyme. Confocal images of *EcRE-lacZ* expression (green) in (**D-F**) dark gut (L3^dark^) or (**H-K**) clear gut (L3^clear^) wing discs expressing (**D**,**H**) RFP only or (**E-F**,**I-K**) EcR^LBD^ in combination with RNAi or overexpression of *EcI* or *Cyp18a1* as indicated. (**G**) As noted in text, *en>Cyp18a1* lethality necessitated the use of *ptc-Gal4* to assess *Cyp18a1* overexpression (oe) effect on EcR^LBD^ and *7xEcRE-GFP*. Yellow arrow in **H** denotes baseline *EcRE-lacZ* activity in the posterior domain of L3^clear^ discs which is partially by rescued by EcI^oe^ or Cyp18a1^RNAi^ in **J** and **K**.

### EcR^LBD^ provides insight into control of physiologic EcR targets

The ability of EcR^LBD^ to distinguish repression and activation phases of *EcRE-lacZ* and *7xEcRE-GFP* provides an opportunity to assess whether physiologic EcR target genes also shift from repression-to-activation as wing cells move through the 3^rd^ instar 20E temporal gradient. *broad complex* (*br*) is an EcR-regulated locus encoding a family of related zinc-finger transcription factors involved in 20E-regulated development of imaginal discs and other larval and pupal tissues [20, 50, 51]. Levels of Br protein are low in early L3 wing nuclei, but increase substantially in response to a 20E pulse ∼5hrs after 3^rd^ instar ecdysis that defines the nutrient-responsive critical weight (CW) checkpoint [52]. Starvation pre-CW leads to developmental arrest as L3 larvae, while post-CW larvae undergo pupation into adults even under low nutrient conditions. To test the effect of EcR^LBD^ on Br expression, *dark, light*, and *clear* L3 *en>EcR*^*LBD*^ wing discs were stained with an antibody that recognizes all Br isoforms (Br^core^) (**Fig. 5A-F**). As described in prior work [52], Br levels are increased substantially in control (*en>RFP*) L3^clear^ wing cells vs L3^dark^ cells (**Fig. 5A,C,E**). Expression of EcR^LBD^ (*en*>*EcR*^*LBD*^) elevates Br levels in L3^dark^ and L3^light^ discs cells relative to control anterior cells (**Fig. 5A-D**) but does not induce Br beyond the high levels already present in L3^clear^ discs (**Fig. 5E-F)**. An identical result is observed when *ptc-Gal4* is substituted for *en-Gal4* **(Fig. S5A-D**). This inability of EcR^LBD^ to induce ectopic Br protein in late-L3 cells is consistent with previous analysis of EcR regulation of a *broad* reporter construct [30] and indicates either that Br levels are already maximal in late-L3 wing cells and cannot be induced further, or that Br repression in late-L3 cells is mediated by a distinct mechanism. A candidate for this role is the NuRD complex component Mi-2 [53], a chromodomain helicase that limits induction of EcR target genes by 20E-induced binding to the AF2 region [54]. A second major conclusion of these data is that LBD-dependent expression of Br occurs primarily by repression, rather than activation. Given that the minimal *EcRE-lacZ* and *7xEcRE-GFP* enhancers are sensitive to both LBD-dependent repression and activation (see **Fig. 2**), it appears that intact EcR-target loci can be sensitive to one or both mechanisms.

**Figure 5.**
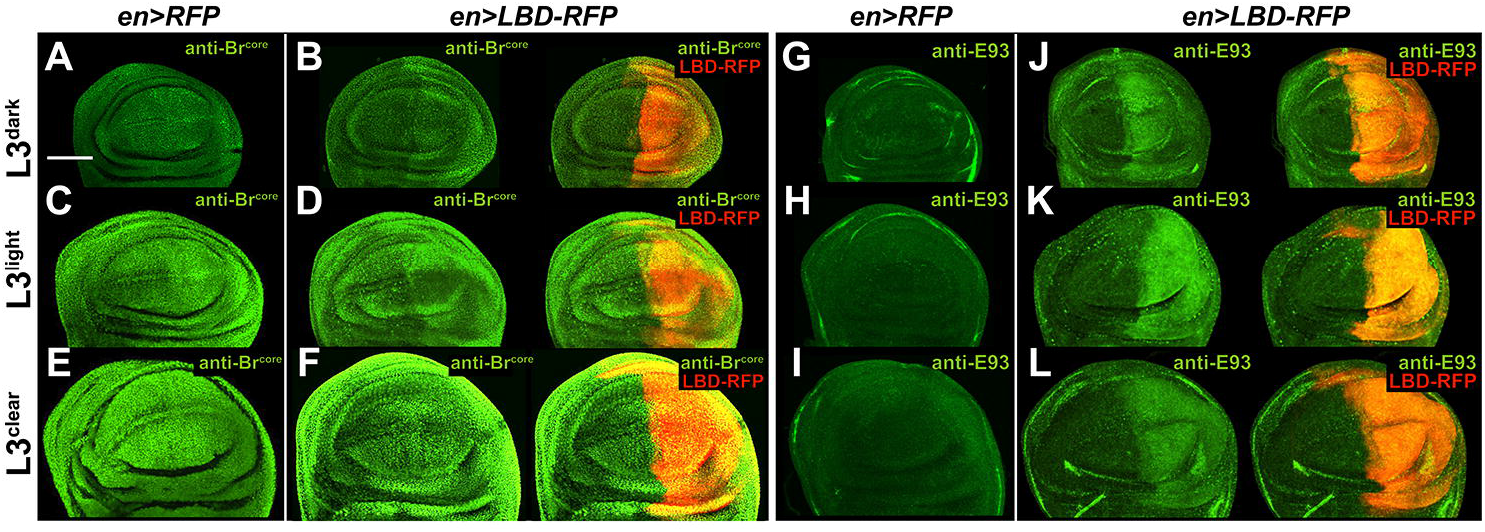
EcR^LBD^ derepresses endogenous EcR targets. Expression of the EcR-induced factors (**A-F**) Broad (Br) and (**G-L**) E93 in control *en>RFP* (**A**,**C**,**E** and **G**,**H**,**I**) or *en>EcR*^*LBD*^ (**B**,**D**,**F** and **J**,**K**,**L**) *dark, light* and *clear* L3 wing discs. Anti-Br^core^ (all Br isoforms) and anti-E93 signals are in green. Red denotes RFP in control panels, and the EcR^LBD^:RFP fusion in all other panels. Note derepression of Br^core^ in the *dark*/*light* but not *clear* timepoints, while E93 is induced at all timepoints.

To extend these analyses to other EcR targets in larval wing disc cells, *EcR*^*LBD*^ was tested for effects on the ecdysone-induced transcription factor E93 (**Fig. 5G-L**) and *ftz-f1, krh-1, tai* and *neb* (**Fig. S6**), which represent candidate EcR targets based on chromatin-occupancy data [30, 46]. *E93* is normally induced in the prepupal-to-pupal stage [55, 56], but EcR^LBD^ induced precocious E93 protein expression in third instar wing discs (**Fig. 5G-L**). A similar derepression is observed for *ftz-f1-lacZ* (**Fig. S4**), which reports expression of the *ftz-f1* transcription factor. *ftz-f1* is part of the genetic hierarchy that controls early metamorphosis downstream of EcR [57], and its expression at the early pupal-to-pupal transition normally coincides with a drop in 20E levels driven by EcR-dependent induction of *Cyp18a1* [35]. Thus, EcR^LBD^ appears to bypass this 20E-EcR-Cyp18a1 feedback mechanism and trigger ectopic expression of *ftz-f1* in larval cells. The *tai-lacZ, krh1-lacZ* and *neb-lacZ* transcriptional reporters are also precociously induced by EcR^LBD^ in L3 wing cells (**Fig. S6**). As with Br, these patterns of reporter induction indicate that the EcR-LBD provides primarily repressive inputs on these genes over developmental time points examined in this study.

### Role of the Smr-EcR interaction in EcR^LBD^ repression

The ability of EcR^LBD^ to modulate expression of EcR target genes and disrupt EcR-regulated processes is consistent with sequestration of EcR co-regulators, including 20E (see **Fig. 4B**). The EcR transcriptional co-regulators Taiman (Tai, [26, 58]) and SMRT-related EcR-interacting protein (Smr, [36]) interact via the AF2 region, and are thus candidates to be bound by EcR^LBD^. Given the strong evidence that LBD derepresses multiple EcR reporters and targets in early/mid-L3 wing cells, attention focused was on the role of the Smr repressor in EcR^LBD^ phenotypes.

Smr is a Myb-SANT domain repressor homologous to the mammalian protein NCoR1 (nuclear receptor corepressor-1, [reviewed in 59]), and is the most well-studied EcR repressor in *Drosophila*. Smr represses EcR activity in diverse tissues, including wing discs [36, 60], where a *Smr-Gal4* trap line (*Smr*^*NP7257*^; Kyoto #105406) driving a CD8:GFP fusion is active throughout the epithelium, with rows of cells with higher Gal4 activity flanking the D:V and A:P boundaries (**Fig. 7B**). To test whether EcR^LBD^ derepression of EcR targets requires Smr binding, the *UAS-EcR*^*LBD*^ system was modified to encode a version of the LBD fragment carrying an Ala^483^ to Thr (A^483^T) mutation in the AF2 region that blocks interaction with Smr [36]. Expression of this Smr-binding mutant form of LDB in the wing posterior domain (*en>EcR*^*LBD-A483T*^) fails to upregulate *EcRE-lacZ* in L3^dark^ cells but mildly represses this reporter in L3^clear^ cells (**Fig. 6A-B**), indicating that the A^483^T mutation abrogates EcR^LBD^ derepression of a simple *EcRE*-based reporter. EcR^LBD-A483T^ has also lost the ability to induce Br^core^ protein in mid-L3 wing cells (**Fig. 6C-D**). This loss of Br^core^ induction by EcR^LBD-A483T^ vs. EcR^LBD^ is also evident using the *patched-Gal4* driver in L3^dark^ discs (**Fig. S5E-F**). These data suggest that EcR recruits Smr to represses Br in pre-CW wing cells but not in post-CW wing cells. EcR^LBD-A483T^ is also significantly impaired in induction of *tai-lacZ* and *krh1-lacZ* but has a similar effect on *neb-lacZ* (**Fig. S6**), suggesting that *neb* may be regulated by a Smr-independent mechanism.

**Figure 6.**
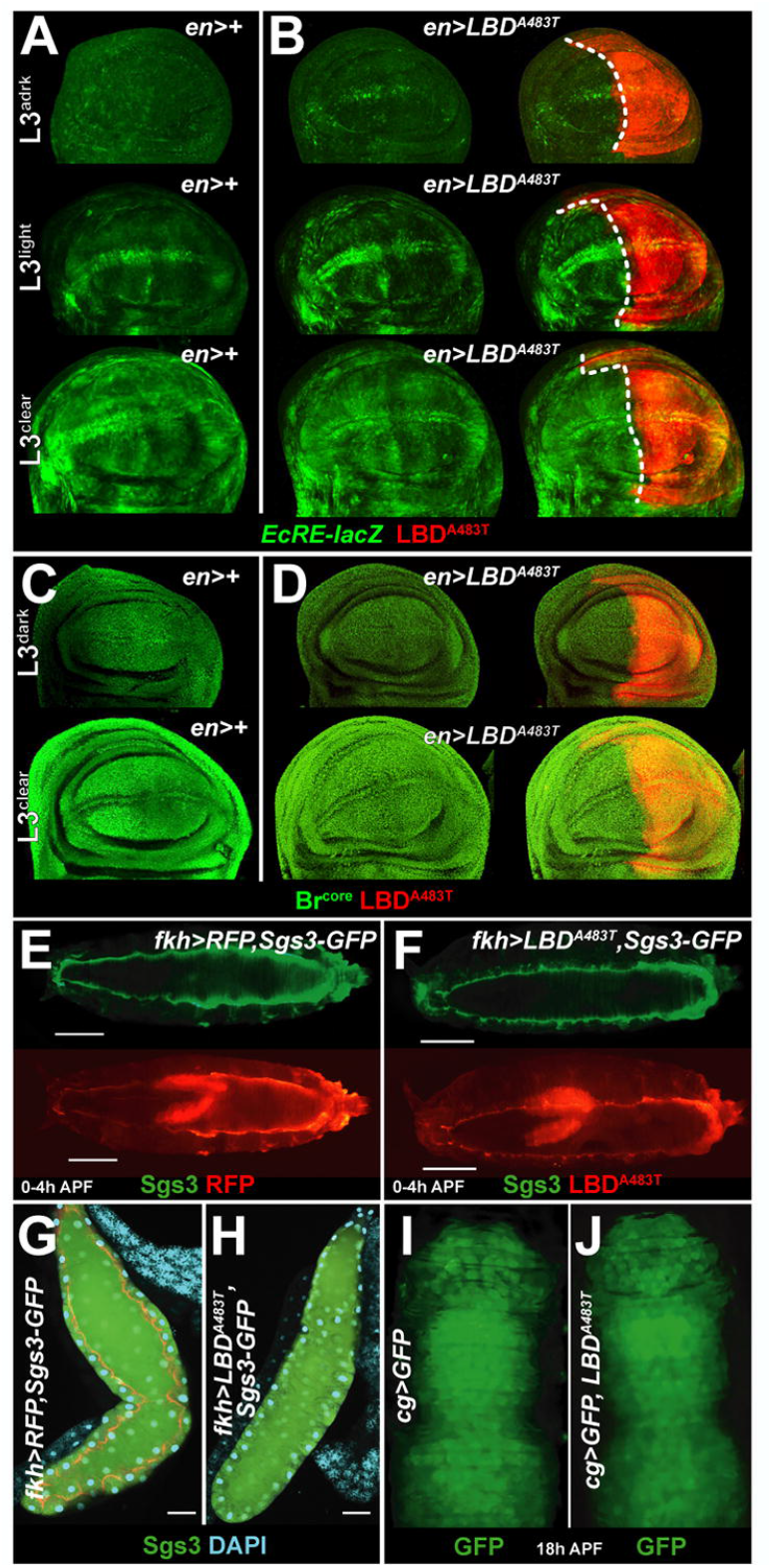
EcR^A483T^ is deficient in repression and activation. Confocal images of *EcRE-lacZ* activity (green) in (**A**) control *en>+* or (**B**) *en>EcR*^*LBD-A483T*^ wing discs collected from *dark, light*, or *clear* L3 larvae. Similar images of Br^core^ signal (green) in (**C**) control *en>+* or (**D**) *en>EcR*^*LBD-A483T*^ discs from *dark* or *clear* L3 larvae. Red indicates the EcR^LBD-A483T^:RFP fusion protein. Note the inability of EcR^LBD-A483T^ to derepress *EcRE-lacZ* in L3^dark^ young discs and fully repress this reporter in older L3^clear^ discs (compare to **panels 2D-F**). Images of (**E**,**G**) control or (**F**,**H**) EcR^LBD-A483T^ expressing 0-4h pupae using the *fkh-Gal4* driver and *Sgs3-GFP* reporter to track glue protein (green). Note the failure of EcR^LBD-A483T^ to block Sgs3 secretion or expectoration. (**I-J**) EcR^LBD-A483T^ also fails to prevent FB migration into the pupal head at 18h APF

**Figure 7.**
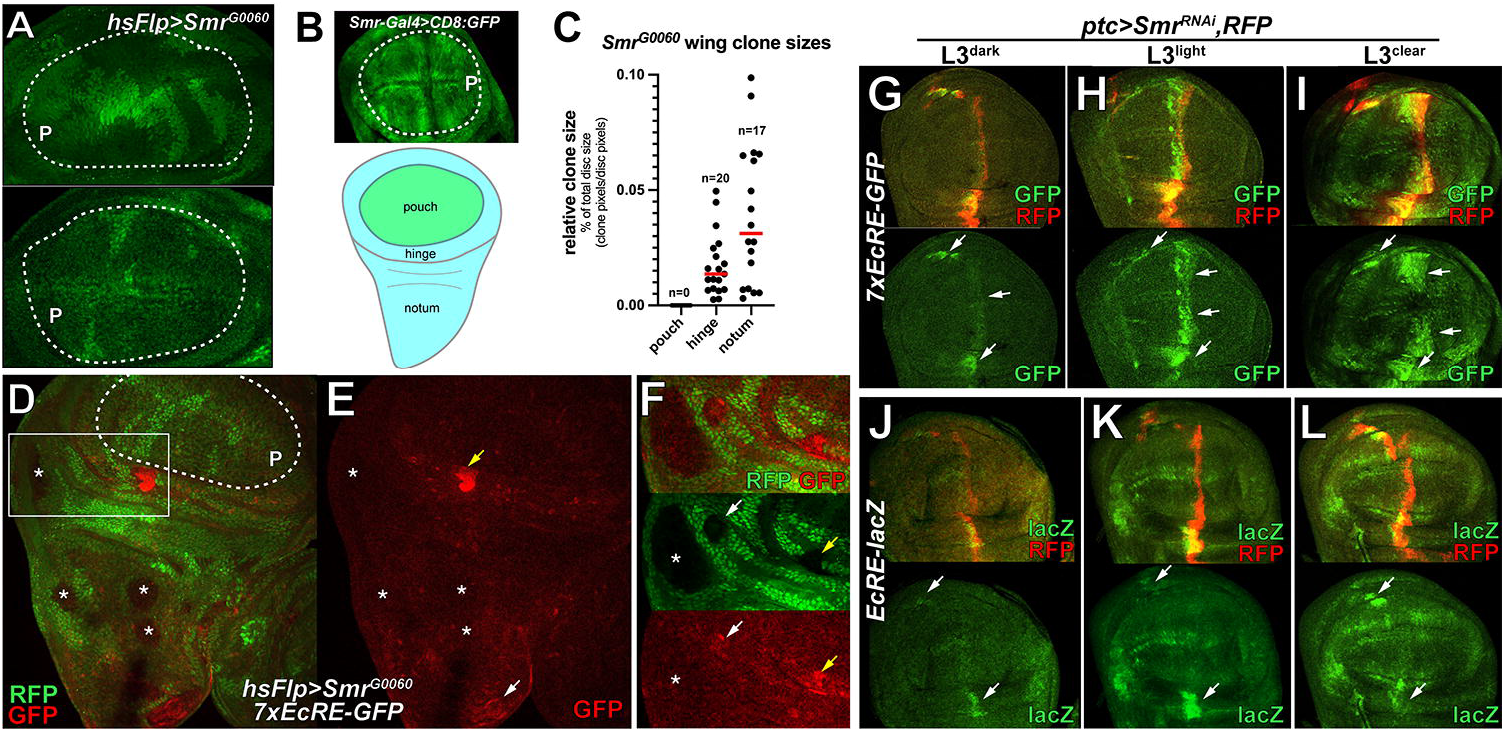
Spatial variation of *EcRE*-reporter repression by Smr. (**A**) Confocal images of wing pouch regions of heat-shock induced *hsFLP;Smr*^*G0060*^ L3 larvae. Note the surviving control twinspots (2x GFP, bright green) but lack of paired *Smr*^*G0060*^ clones (lacking GFP, black). (**B**) Pouch and hinge expression of a *Smr-Gal4* line (*P{GawB}Smr*^*7257*^) driving *UAS-CD8:GFP* (green). (**C**) Dot plot analysis of two-dimensional *Smr*^*G0060*^ clone size in the pouch, hinge, and notum (see cartoon). Clone sizes are normalized to total disc size. Numbers of clones (n) for each condition are indicated. Red bars denote the mean clone size. (**D-E**) Effect of *Smr*^*G0060*^ clones (marked by absence of RFP; false colored green) on *7xEcRE-GFP* (false colored red) in an L3 wing disc. Note absence of surviving clones in the pouch (dotted line). Asterisks denote *Smr*^*G0060*^ clones in the hinge and notum that elevate *7xEcRE-GFP* weakly or not at all. Yellow arrow in (**E)** marks a medial hinge *Smr*^*G0060*^ clone that strongly activates *7xEcRE-GFP*; white arrow marks a clone at the far dorsal tip of the disc that weakly activates *7xEcRE-GFP*. Inset in (**F**) shows a more basal optical slice of *7xEcRE-GFP* expression (boxed in **D**) in the same medial hinge clone in (yellow arrow in **E**), and a small hinge clone that weakly activates *7xEcRE-GFP* (white arrow), and a larger notum clone that has no effect on *7xEcRE-GFP* (asterisk). (**G-L**) Images of *dark/light/clear* timed wing discs showing temporal and spatial variation in *ptc>Smr*^*RNAi*^ derepression of the (**G-I**) *7xEcRE-GFP* (green) or (**J-L**) *EcRE-lacZ* (green) reporters. RFP denotes the *ptc* domain (*ptc>RFP*). Arrows indicate areas of stronger reporter induction in each panel.

At the organismal level, the *A483T* mutation eliminates the ability of EcR^LBD^ to block Sgs3-GFP secretion into the larval SG lumen and WPP stage expectoration of glue proteins (**Fig. 6E-H**). These data imply either that Smr is required to coordinate 20E-induced Sgs3 production, secretion, and expectoration, or that the A^483^T mutation also disrupts binding of an unidentified co-activator that promotes these processes. A direct test of these models by RNAi knockdown of *Smr* in larval SGs (*fkh>Smr*^*IR*^) reveals that Smr is required for Sgs3-GFP secretion (see *Smr*^*RNAi*^ data in **Fig. S1E**), perhaps by coordinating a shift in EcR-regulated gene expression from glue protein production to secretion/expectoration. Relative to EcR^LBD^, EcR^LBD-A483T^ has also lost the ability to block FB migration into the pupal head (**Fig. 6I-J** vs. **Fig. 1C**). As in the SGs, this result seems to imply that a balance of EcR repression and activation is required for proper timing of EcR-regulated developmental processes.

To independently assess the role of Smr in repressing EcR activity in larval wing cells, a strong loss-of-function *Smr* allele (*Smr*^*G0060*^,*FRT19A*) was used to create homozygous mutant *Smr* clones using the *hs-Flp:FRT* system. Precise excision of the P-element in *Smr*^*G0060*^ rescues lethality [60], indicating that it is an appropriate genetic background for phenotypic studies. Upon dissection, it was noted that the presence of *Smr*^*G0060*^ clones enhances wing disc folding, especially in the hinge region, with clones optimally visible in different Z-planes. At 96hrs post heat-shock (hs), L3 wing discs lacked *Smr*^*G0060*^ clones in the wing pouch (P), despite evidence of surviving GFP-positive twinspots throughout the pouch (**Fig. 7A-B**). This apparent cell-lethal effect of *Smr*^*G0060*^ in the pouch contrasts with the effect of *Smr* loss in the hinge and notum, where *Smr*^*G0060*^ clones are consistently recovered following hs-Flp induction (**Figs. 7C-D** and **S7A**). *Smr*^*G0060*^ notum clones are on average 2-3 times larger than hinge clones (**Fig. 7C**), indicating that proliferative effects of Smr loss vary by region in the wing epithelium.

To test the effect of *Smr*^*G0060*^ on EcR activity in different wing regions, *Smr*^*G0060*^ clones were generated in the background of the *7XEcRE-GFP* transgene using *hsFlp:FRT* (**Figs. 7** and **S7A**). The absence of *Smr*^*G0060*^ clones in the pouch precluded analysis of *7xEcRE-GFP* in this region (e.g., area inside dotted line in **Fig. 7D**) but clones located in the hinge and notum displayed location-dependent *7xEcRE-GFP* activation. *Smr*^*G0060*^ hinge clones located along the medial axis and immediately dorsal to the pouch showed strong activated *7xEcRE-GFP* (yellow arrow, **Fig. 7E,F**) while *7xEcRE-GFP* was more weakly activated in more lateral hinge clones located further from the pouch. A magnified view of the boxed region in **Fig. 7D** shows a basal optical slice containing the same medial clone as in **Fig. 7E** with high *7xEcRE-GFP* (**Fig. 7F**, yellow arrow) and a lateral clone with lower *7xEcRE-GFP* (white arrow). *Smr*^*G0060*^ clones in the notum did not activate *7xEcRE-GFP* (asterisks in **Fig. 7D-F**), except for occasional *Smr*^*G0060*^ clones located near the dorsal tip the notum (white arrow, **Fig. 7E**). In sum, these data indicate that Smr represses transcriptional output of the *7xEcRE-GFP* reporter in specific regions of the wing disc, with medial hinge cells immediately below the pouch showing the strongest link between EcR activity and Smr loss, and that the effects of Smr on clone size varies independently of the effect on *7xEcRE-GFP*, with Smr loss associated with clone expansion in the notum and elimination in the pouch.

To bypass the cell-lethal effect of *Smr*^*G0060*^ in wing pouch cells, *patched-Gal4* was used to drive a *Smr* RNAi transgene (BL# 27068) in the background of *7xEcRE-GFP* or *EcRE-lacZ* and timed by the blue-dye method [48]. In L3^dark^ discs, *ptc>Smr*^*RNAi*^ elevated *7xEcRE-GFP* in dorsal and ventral hinge cells located at the A:P boundary, and in midline cells of the pouch (arrows, **Fig. 7G**). This pattern was intensified in L3^light^ discs and resolved into two bright wedges of *7xEcRE-GFP* activity oriented along the dorsoventral axis of the wing pouch (**Fig. 7H-I**). For comparison, knockdown of a second EcR-associated repressor, the Cop9 signalosome subunit-2 (CSN2; also known as *alien* [61]), produces mild induction of *7xEcRE-GFP* (**Fig. S5D**), suggesting a greater requirement for Smr than CSN2 in repressing of this reporter in L3 wing cells. In contrast to *7xEcER-GFP*, the *EcRE-lacZ* reporter was only weakly activated by *ptc>Smr*^*RNAi*^ in L3^dark^ ventral midline hinge cells, but strongly activated in dorsal hinge cells from L3^dark^ and L3^light^ discs (**Fig. 7J-L**). Each of these patterns were also observed in *en>Smr*^*RNAi*^ discs (**Fig. S5B-E**), which further confirmed that Smr strongly represses *7xEcRE-GFP* in dorsal and ventral hinge cells located along the AP boundary in L3^dark^ and L3^light^ discs, but strongly represses *EcRE-lacZ* only in the dorsal group in L3^light^ discs (circled in **Fig. S5B-C**,**F**). In sum, these data reveal that Smr represses EcR activity in wing cells throughout the L3 period, even after the CW 20E pulse and accompanying repression-to-activation switch detected with EcR^LBD^ (see **Fig. 2**). This finding suggests that a 20E pulse can tip inputs in favor of activation on a given enhancer, but that Smr repression continues to limit the extent of this activation on EcR targets after the mid-L3 20E, especially in dorsoventral midline cells located in the hinge. EcR^LBD^ appears capable of blocking both Smr-mediated repression, as indicated by loss of de-repression activity of EcR^A483T^, and 20E-driven activation, with each effect likely mediated by sponging of key cofactors by the AF2 and LBD regions.

## Discussion

Here we employ a Gal4/UAS expressed EcR^LBD^ ‘sponge’ that titrates EcR co-regulators and permits assessment of expression of individual target genes *in vivo* without disrupting endogenous EcR bound at genomic sites. Within 3^rd^ instar wing discs, these experiments reveal a dramatic temporal switch from EcR-mediated repression to activation that occurs in a narrow temporal window that precedes the large pulse of 20E at the larval-to-pupal transition. Molecular data link the EcR^LBD^ repressive role to physical binding to the activating ligand 20E, and its activating role to sequestering the Smr co-repressor. EcR^LBD^ derepression of heterologous EcR reporters in young L3 wing discs is genetically insensitive to depletion of the *Oatp74D* ecdysone importer EcI, suggesting that relief of repression can occur without concomitant activation by 20E-liganded EcR. Reciprocally, EcR^LBD^ repression of these same reporters in late L3 wing cells is partially rescued by depletion of *Cyp18a1*, which inactivates 20E, indicating that EcR^LBD^ repression is overcome by elevating available intracellular 20E. Consistent with these models, a version of EcR^LBD^ carrying the Ala^483^Thr mutation, which blocks interaction with the Smrter (Smr) corepressor [36], is defective in induction of EcR targets in young L3 wing cells. including the well-characterized EcR target Broad (Br). Unexpectedly, EcR^LBD- A483T^ is also defective in repression of endogenous EcR activity in late L3 wing cells, suggesting that 20E-induced eviction of Smr may result in binding of coactivators to EcR using an interaction interface shared by Smr. This data suggests that *in vivo* phenotypes associated with the *EcR*^*A483T*^ allele are caused by a lack of repression by Smr, but perhaps also a defect in LBD/AF2 dependent activation. Direct tests of *Smr* roles wing disc cells using clonal mosaics and RNAi confirms a requirement to repress EcR transcriptional activity, but also reveals unexpected regional differences in *Smr* mutant phenotypes, with the greatest effect on EcR activity observed in medial hinge cells and dramatically enhanced expansion of *Smr* mutant clones in the notum relative to age-matched clones in the pouch. In sum, these studies reveal a temporal switch between EcR repression and activation in L3 larval wing cells that is sensitive to genetic manipulation of local 20E levels and connect these mechanisms to EcR-Smr dependent repression of EcR targets. This evidence indicates that repression and derepression may be a common mode of transcriptional regulation exerted by 20E-EcR signaling during the third instar. Finally, these data validate the EcR^LBD^ fragment as an effective tool to assess the relative contribution of EcR repressors and activators to temporal control of individual EcR targets *in vivo* and provide a proof-of-principle that transcriptional effects of other *EcR* mutations that lie in LBD/AF2 region can be effectively modelled in the EcR^LBD^ system.

The EcR repression-to-activation switch in wing disc cells occurs between 120-144h AEL, which precedes the large pulse of 20E that occurs as larvae reach the end of L3 and enter pupariation [62]. This pulse of 20E has long been presumed to drive EcR activity, so it is notable that two distinct *EcRE*-based reporters detect a clear repression-to-activation switch in response to comparatively small peaks of 20E in mid-L3. One source of this sensitivity could be a lower threshold for 20E activation of EcR in wing cells, perhaps because of more permissive chromatin environments or cooperating transcription factors bound on the same loci [63], or an elevated ability to uptake Ec via importers, e.g., Oatp74D and Oatp33Eb [5, 64]. In this regard, it is notable that past studies have measured 20E levels in whole animal lysates [e.g., 65], which may not accurately reflect intracellular hormone levels. Somewhat unexpectedly, although both

*EcRE-lacZ* and *7xEcRE-GFP* reporters are both derepressed in L3^dark^ wing cells and repressed in L3^clear^ wing cells, reporters of EcR-bound promoters tested in this study, including *br, E93, ftz-f1, tai*, and *neb* [30], all respond to *EcR*^*LBD*^ expression by precocious activation with no evidence of repression in late L3 wing cells. This gap could reflect the relatively small number of EcR targets tested in the current study. However, it also supports recent evidence that a significant fraction of physiologic EcR targets in L3 wing cells are only bound by EcR when they are repressed, including *br* [30]. Other genes are bound only by EcR in early pupal wing cells, while others are bound in both pre- and post-pupation wing cells. The ability of EcR^LBD^ to block activation of *EcRE*-reporters in late L3 wing cells argues that EcR^LBD^ is ideal system to test the hypothesis that EcR occupancy on genes in late-L3 and early pupae stages predict a requirement for EcR-20E to activate their transcription.

Overexpression of dominant *EcR* alleles (e.g., *EcR*^*F645A*^ [38]) or *EcR*^*RNAi*^ will either displace or remove endogenous EcR, while EcR^LBD^ will only compete away co-regulators bound to the hinge and LBD/AF2 domains of endogenous EcR. Interactions with cofactors that occur through other domains, such as the ligand independent AF1 domain [14, 66], are predicted to remain intact in cells that express EcR^LBD^. This key difference may explain why EcR^LBD^ phenocopies some *EcR* loss-of-function phenotypes [33] but not others. For example, *EcR*^*F645A*^ and *EcR*^*RNAi*^ each disrupt production, secretion, and expectoration of glue proteins, but *EcR*^*LBD*^ only disrupts secretion and expectoration. Similarly, *EcR*^*F645A*^ and *EcR*^*RNAi*^ each block FB rearrangement and migration into head, while *EcR*^*LBD*^ expression only blocks FB migration. These phenotypic differences could highlight processes that require AF1-mediated interactions or cooperative effects between AF1 and AF2, which can work together to regulate hormone receptor effects in other organisms [67-72]. Indeed, future comparisons between *EcR*^*LBD*^ and *EcR*^*AF1*^ transgenes could be an effective way to dissect roles of AF1 in 20E-EcR driven phenotypes in developing tissues. Coupling these functional studies with proteomic analysis of factors that bind EcR^LBD^ and EcR^AF1^ in intact tissues (e.g., larval discs) would be an effective way to link specific EcR-cofactor interactions to regulatory effects on specific downstream targets. As most coregulator interactions mediated through the AF1 and AF2 domains are conserved between *Drosophila* and mammalian NHRs, these insights into EcR-cofactor interactions could also provide insight to the functions of mammalian NHRs [73, 74].

## Acknowledgements

We thank the Emory Integrated Genomics Core and the Emory Integrated Cellular Imaging Core for their expertise and assistance. We also thank R. Evans, N. Yamanaka, D. McKay, A. Bashirullah, V. Henrich, the Developmental Studies Hybridoma Band (DSHB), Bloomington *Drosophila* Stock Center (BDSC), the Harvard Transgenic RNAi Project (TRiP), and the Kyoto Stock Center for antibodies and stocks. We thank ACO, ACO, and LMO for patience and support. We also thank members of the Moberg and Robinson laboratories for their helpful discussion. This work was funded by NIH GMxxxxx (to KHM) and NIH K12-GM000680 (to JWO) and a Mark R. Hudgens Postdoctoral Fellowship in Cancer Research from the American Cancer Society and Scott Hudgens Family Foundation (ACS PF DDC-132078 to JWO).

## Author Contributions

Conceived and designed the experiments: JWO, KHM. Performed the experiments: JWO, CS, DT, OS, MT, DS. Analyzed the data: JWO, KHM. Wrote the paper: JWO. Edited the manuscript: JWO, KHM.

## MATERIALS AND METHODS

### *Drosophila* husbandry

All *Drosophila* stocks were maintained at 25°C unless otherwise noted. Larval staging was performed using the blue food method as described in [48]. *7xEcRE-GFP, sgs3-Gal4;sgs3-GFP* and *UAS-EcI* were gifts of V. Henrich, A. Bashirullah, and N. Yamanaka, respectively. Remaining stocks were obtained from the Bloomington Drosophila Stock Center (BDSC) or the Kyoto Stock Center: *fkh-Gal4* (BL#78061), *sgs3-GFP* (BL#5884), *cg-Gal4* (BL#7011), *Act-5c-Gal4* (BL#25374), *UAS-EcR-RNAi* (BL#9327), *UAS EcI-RNAi* (Kyoto#7571R-1), *UAS-EcRF*^*645A*^ (BL#9452), *UAS-Cyp18a1* (BL#19262), *EcRE-lacZ* (BL#4517), *ptc-Gal4* (BL#2017), *UAS-Cyp18a1-RNAi* (BL#64923), *kr-h1-lacZ* (BL#10381), *tai-lacZ* (BL#11045), *neb-lacZ* (BL#10391), *ftz-f1-lacZ* (BL#11598), *ubi>mRFP,hsFlp,FRT19A* (BL#31418), *Smr*^*G0060*^ (BL#5596), *Smr*^*BG0648*^ (BL#13116), *Smr-Gal4* (BL#76743), *UAS-RFP* (BL#30556), *UAS-CD8:GFP* (BL#32184), and *UAS-Smr-RNAi* (BL#27068).

### *UAS-EcR*^*LBD*^ and *UAS-EcR*^*A483T*^ transgenic lines

RedStinger (nlsRFP) coding sequence was PCR amplified from *IRER-nlsRFP* genomic DNA [75] with primers containing 5’ and 3’ Not1 and Xba1 restriction sites. Not1/Xba1 digested pUAST and PCR product were ligated at 16ºC. The resulting *pUAST-nlsRED* plasmid was sequence verified before being subsequently digested with EcoR1/Not1 to allow for the insertion of DNA encoding the EcR^LBD^, hinge region and F-domain (amino acids 330-878) which were PCR amplified from *hs-EcR*.*LBD* genomic DNA with EcoR1/Not1 restriction sites added [6, 33]. The completed *pUAST-EcR*^*LBD*^*-RFP* plasmid was sequence verified. The Emory Integrated Genomics Core introduced the A^483^T single point mutation using standard practices to generate *pUAST-EcR*^*A483T*^. Transgenic fly services were provided by BestGene, Inc (USA).

## Analysis of LBD phenotypes

### FB migration

GFP was expressed in the larval fat body (FB) by intercrossing *cg-Gal4* and *UAS-CD8:GFP* lines. White prepupae were identified and collected at 18h or 24h after pupariation then transferred to a glass slide for imaging. Images were taken on a Leica MC170 HD camera at 25X magnification (2.5x objective x 10x eyepiece) using the Leica Application Suite and a Zeiss AttoArc HBO 100W lamp.

### Glue protein dynamics

To measure glue protein secretion into the lumen, late (clear gut) 3^rd^ instar *sgs3-GFP* larvae of the indicated genotypes were dissected, fixed in 4% paraformaldehyde for 30min, washed 3x 5min in 1x phosphate buffered saline (PBS), permeabilized with 1xPBS+0.3% Triton-X 100, and stained with DAPI (Sigma Aldrich #D8417) for 10min. To assess glue expectoration, *sgs3-GFP* pupae (4h APF) were imaged through their pupal case for GFP localization and intensity. Sgs3-GFP was scored as ‘retained’ if GFP signal could be seen within the SGs.

### Mouth hook retention

To assess mouth hook retention, 4h collections of eggs laid on yeasted grape juice plates at 25ºC were aged for an additional 24h. Early first-instar larvae of the correct genotype were collected and transferred to standard food. Larvae were then collected at 24h and 48h later to analyze mouth hooks following L2 and L2 ecdysis.

### Developmental delay

4h old eggs laid on yeasted grape juice plates at 25ºC were aged for an additional 24h, and first instar larvae of the correct genotypes were collected and transferred to standard food and dissected at the hours indicated.

### Immunohistochemistry

Larval tissue was processed as previously described [63, 76]. Briefly, discs were dissected in 1xPBS, fixed 20min in 4% paraformaldehyde at room temperature, rinsed 3x in 1x PBS, then permeabilized for 30min at RT in 1xPBS+0.3% Triton X-100. Samples were then blocked for 30min in in 1xPBS+10%normal goat serum (NGS; Jackson Immunologicals). Samples were then resuspended in primary antibody in 1xPBS+0.1% Triton X-100+10%NGS and incubated overnight at 4ºC. Following 5x 5min washes in 1xPBS+0.1% Triton X-100, samples were then incubated for 45min at room temperature in secondary antibody resuspended according to manufacturer’s instructions in 1xPBS+0.1% Triton-X+10% NGS. After 5x 5min washes in 1xPBS+0.1% Triton X-100 discs were mounted in *n*-propyl gallate (4% w/v in glycerol). The following antibodies were used: mouse anti-Broad core (1:100; Developmental Studies Hybridoma Bank #25E9.D7), mouse anti-β-Gal (1:500; Promega Z3781), mouse anti-E93 (1:5000; gift of D. McKay). Nuclei were stained with DAPI for 10min (Sigma Aldrich #D8417). Secondary antibodies were goat-anti rabbit Alexa-488 (1:100, Jackson ImmunoResearch), goat anti-mouse Cy3 (1:50; Jackson ImmunoResearch), and goat-anti-mouse Alexa-647 (1:50; Jackson ImmunoResearch). Wing discs were imaged on a Nikon A1R HD25 confocal system. Images were analyzed in Fiji and processed in Photoshop 2021.

### Immunoprecipitation

4h collections of embryos were aged for an additional 24h yeasted grape juice plates at 25ºC. Early first instar larvae were then transferred to standard food and aged for 48h prior to collection on ice and washing in 1xPBS to remove food particles. Larvae were then lysed with a mortar and pestle in RIPA buffer (10 mM Tris/Cl pH 7.5, 150 mM NaCl, 0.5 mM EDTA, 0.1 % SDS, 1 % TritonTM X-100, 1 % deoxycholate) supplemented with cOmplete Protease Inhibitor Cocktail (Roche; #11697498001) at 4ºC for 5min and then rocked for n additional 30min to ensure complete lysis. Samples were then spun for 15min at 16,000g and supernatant stored at -80 ºC. Samples pooled as needed for immunoprecipitation with RFP-Trap Agarose beads or control beads (Chromotek, #RTA-20; #BAB-20). Agarose beads were equilibrated and 125µL of bead slurry was added per 500 larval lysates. Lysate and bead mixture was tumbled end over end for 1 hour at 4ºC. Beads were then sedimented and washed before moving onto analysis for either immunoblot or ELISA.

### Enzyme-linked immunosorbent assay (ELISA)

To extract 20E bound to the EcR^LBD^, RFP-Trap or binding control agarose beads, immunoprecipitated material (see above) was extracted with 0.5mL methanol twice before a final extraction in 0.5mL ethanol. The samples were dried with a Speedvac concentrator and dissolved in EIA buffer (Cayman Chemical; #501390). The samples were assayed according to the manufacturer’s instructions using synthetic 20E as a standard, and 20E AChE tracer, 20E EIA antiserum, precoated (mouse anti-rabbit IgG) ELISA 96-well strip plate and Ellman’s reagent prepared as instructed. Samples were read on a BioTek platereader at 405nM.

### Nuclear fractionation

To isolate nuclei, larvae were collected on ice and washed as described above. Larvae were then transferred to a clean Dounce homogenizer (Thomas; size AA) and lysed in a hypotonic cell lysis buffer (10mM HEPES, pH 7.6, 1.5mM MgCl_2_, 10mM KCl, 1mM DTT) supplemented with cOmplete Protease Inhibitor Cocktail) for 5 min at 4ºC. Lysates were then transferred to a microcentrifuge tube and spun for 1 min at 60g at 4ºC to pellet the carcass and debris. Supernatant was transferred to a clean microcentrifuge tube and the nuclei were pelleted at 3300g for 10min at 4ºC. The supernatant was discarded, and nuclei were resuspended in 3 volumes of high salt buffer (0.5M NaCl, 10mM HEPES, pH 7.6, 2.5mM MgCl_2_, 0.5mM EDTA, 20% glycerol, 0.1% NP-40, cOmplete Protease Inhibitor Cocktail) and rocked for 30min at 4ºC. The samples were then sonicated and DNA was then digested using DNase (Qiagen, #79254) rotating for 90min at 4ºC. Samples were then diluted with dilution buffer at a 1:2.3 ratio (10mM HEPES, pH 7.6, 1mM DTT, 0.65% NP40, 2.5mM MgCl_2_, 0.5mM EDTA, cOmplete Protease Inhibitor Cocktail) and rocked for 30min at 4ºC. The insoluble nuclear fraction was pelleted at 16,000g for 30min at 4ºC. The soluble nuclear fraction was removed and used for further analysis.

### Immunoblot Analysis

Following IP, 2x Lammeli buffer with 1mM DTT was added to control and RFP-Trap beads, briefly boiled, and eluate run on 4-15% Mini-PROTEAN TGX Stain-Free Precast gels (BioRad; #4568046) and subjected to standard immunoblot techniques. The following antibodies were used for immunoblotting: rabbit-anti-Smrter (1:50, R. Evans), mouse-anti-EcR (1:100; Developmental Studies Hybridoma Bank #DDA2.7), and mouse-anti-lamin (1:100; Developmental Studies Hybridoma Bank #ADL84.12).

### Statistics

Statistical analysis was completed using GraphPad Prism v9.3.1.

