## Supplemental Figs and Legends for "An EcR probe reveals mechanisms of the ecdysone-mediated switch from repression-to-activation on target genes in the larval wing disc"

**Supplemental Figures**

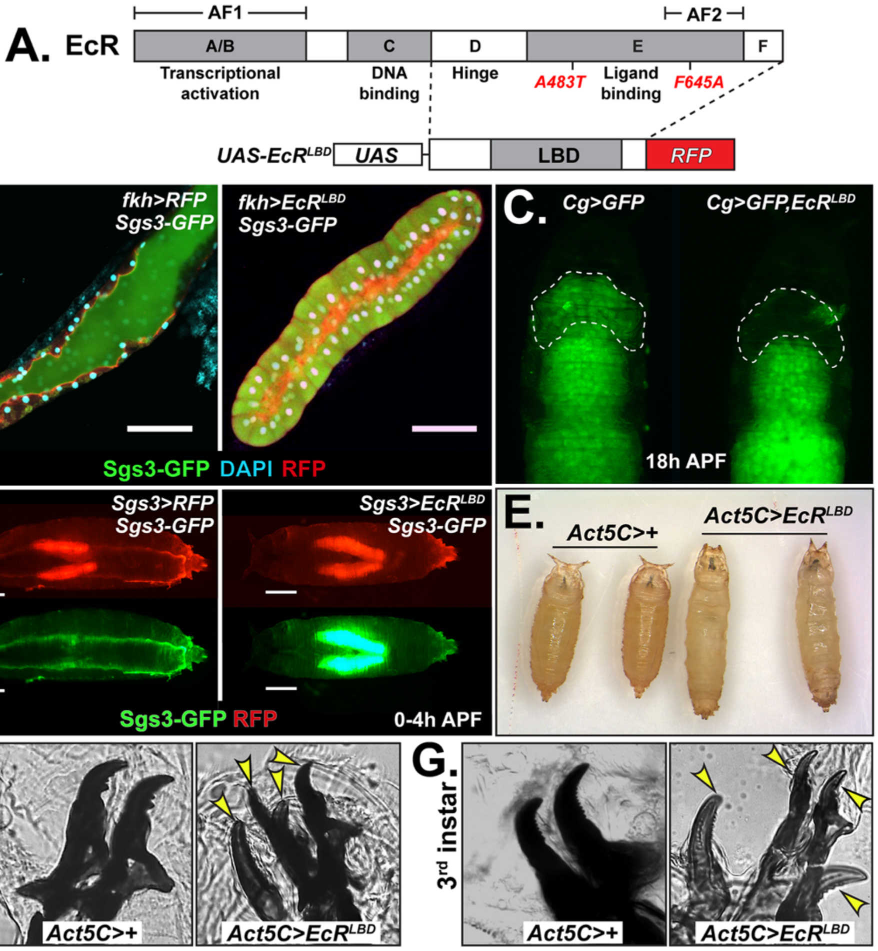


**Supplemental Figure 1. Effects of *EcR*, *Smr* and *Cyp181* transgenes on 20E-EcR regulated processes.** (**A-C**) Compared to *fkh>Sgs3-GFP* control (**B**), expression of $EcI^{RNAi}$, the $EcR^{F645A}$ dominant negative, or $EcR^{RNAi}$ reduces Sgs3-GFP (green) production and luminal secretion in the SGs. (**D**) Compared to *Sgs3>Sgs3-GFP* control, expression of $EcR^{RNAi}$ or $EcR^{F645A}$ prevents Sgs3-GFP (green) expectoration at 0-4h APF. (**E**) Percentage of SGs with Sgs3-GFP retained in control (*fkh>Sgs3-GFP*), $EcR^{LBD}$, $EcR^{RNAi}$, $EcR^{LBD-A483T}$, $Smr^{RNAi}$, $EcR^{F645A}$, and $Cyp18a1^{oe}$ larvae. (**F**) Compared to control (*Cg>GFP*), expression of $EcI^{RNAi,}$ $EcR^{F645A}$, or $EcR^{RNAi}$ prevent the FB (green) remodeling and migration into 18h APF pupal heads (dotted outline). (**G**) Adult survival among F1 progeny ubiquitously overexpressing $EcR^{LBD}$. Chi-square calculated $p$ values are included.

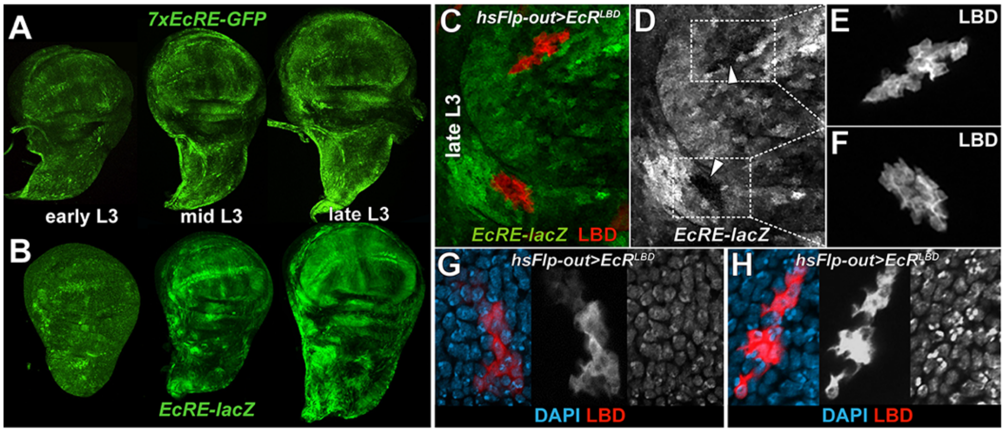


**Supplemental Figure 2. Baseline expression of *EcRE* reporters and EcR$^{LBD}$ localization.** L3 wing discs of the indicated genotype immunostained to detect (**A**) GFP or (**B**) β-galactosidase. (**C-D**) L3$^{clear}$ wing imaginal discs bearing *hsFlp-out* clones of *EcR$^{LBD}$* (red) in the background of *EcRE-lacZ* (green). Panel (**D**) shows a paired grey-scale image of *EcRE-lacZ* activity with arrowheads on EcR$^{LBD}$-expressing clones. (**E,F**) Grey-scale images of EcR$^{LBD}$ expression in clones from panel **C**. (**G,H**) *hsFlp-out* clones of EcR$^{LBD}$ (red) expressing cells in L3$^{clear}$ wing discs co-stained with DAPI (blue).

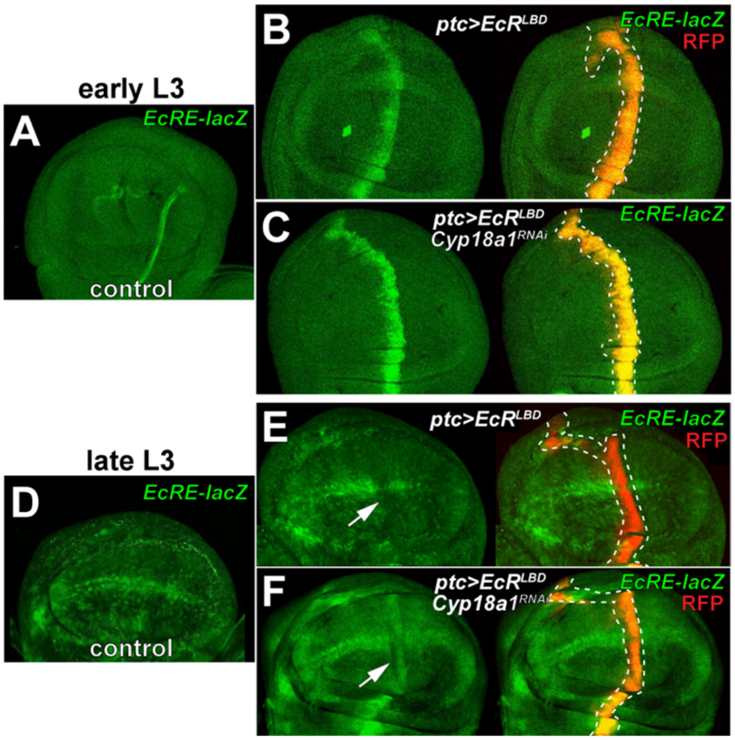


**Supplemental Figure 3. Elevated 20E modulates EcR$^{LBD}$ effects on EcR activity.** (**A-C**) *EcRE-lacZ* activity (green) in early L3 wing discs of the indicated genotypes. Compared to (**A**) *ptc>RFP* control, (**B**) EcR$^{LBD}$ (*ptc>EcRE$^{LBD}$*; red) elevates *EcRE-lacZ* expression, and this is (**C**) further enhanced by *Cyp18a1$^{RNAi}$*. (**D-F**) *EcRE-lacZ* activity (green) in late L3 wing discs of the indicated genotypes. Compared to (**D**) *ptc>RFP* control, (**E**) EcR$^{LBD}$ (*ptc>EcRE$^{LBD}$*; red) represses *EcRE-lacZ* expression, and this is (**F**) partially rescued by *Cyp18a1$^{RNAi}$*.

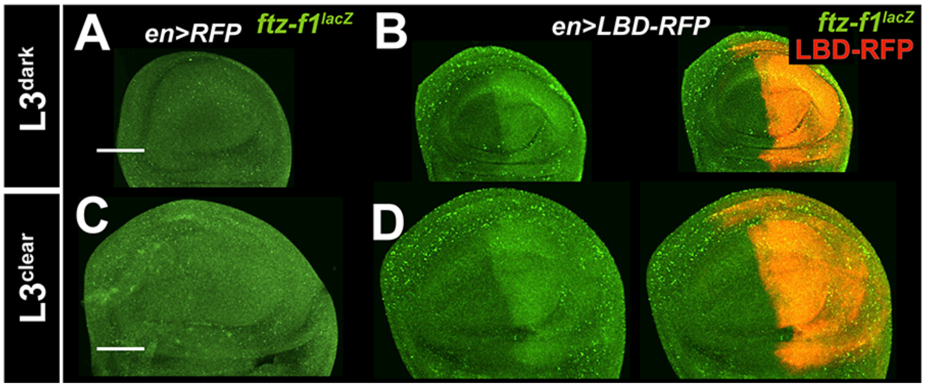


**Supplemental Figure 4. EcR$^{LBD}$ derepresses *ftz-f1* across all L3 timepoints.** Expression of a *ftz-f1-lacZ* transgene (green) in L3$^{dark}$ and L3$^{clear}$ wing discs from (**A,C**) control *en*>+ and (**B,D**) *en>EcR$^{LBD}$* discs (LBD=red).

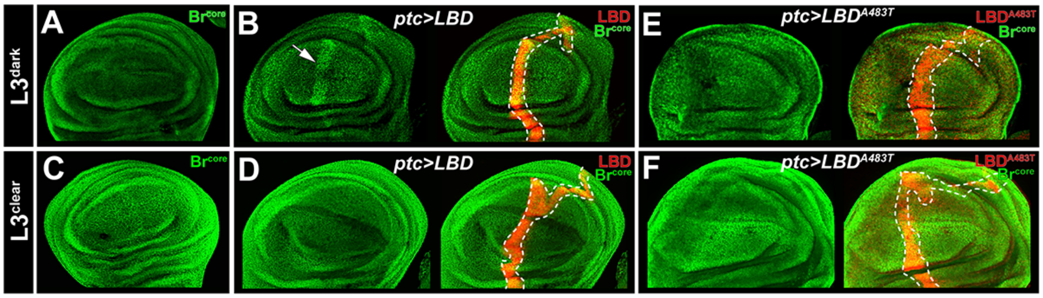


**Supplemental Figure 5. The *EcR$^{A483T}$* mutation disrupts Broad repression**. Expression of Br$^{core}$ isoforms (green) in L3$^{dark}$ and L3$^{clear}$ from (**A,C**) wildtype control, (**B,D**) *ptc>EcR$^{LBD}$* (red) or (**E,F**) *ptc>EcR$^{LBD\text{-}A483T}$* (red) wing discs. Dotted lines outline the LBD:RFP expressing *ptc* domain. Arrow denotes Br$^{core}$ induction only in *ptc>EcR$^{LBD}$* L3$^{dark}$ discs.

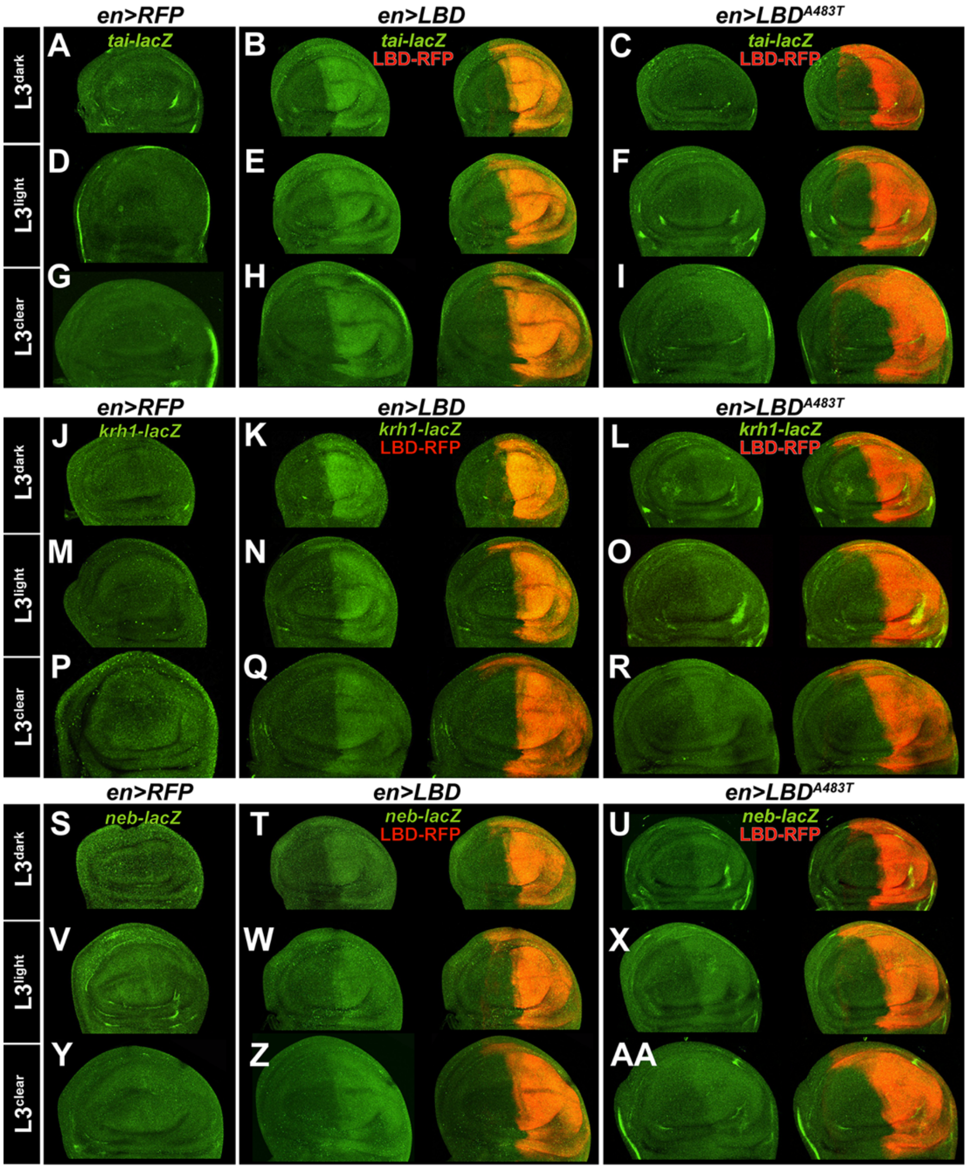


**Supplemental Figure 6**. **The A483T mutation reveals varying levels of Smr repression of EcR-regulated genes.** Expression patterns of the (**A-I**) *tai-lacZ*, (**J-R**) *krh1-lacZ*, and (**S-AA**) *neb-lacZ* reporters in $L3^{dark}$, $L3^{light}$, and $L3^{clear}$ discs of the indicated genotypes. As noted in text, these targets were chosen based on their identification as EcR-bound and regulated genes in wing discs [30]. Note that $EcR^{LBD\text{-}A483T}$ is impaired in induction of *krh1* and *tai*, but not *neb*.

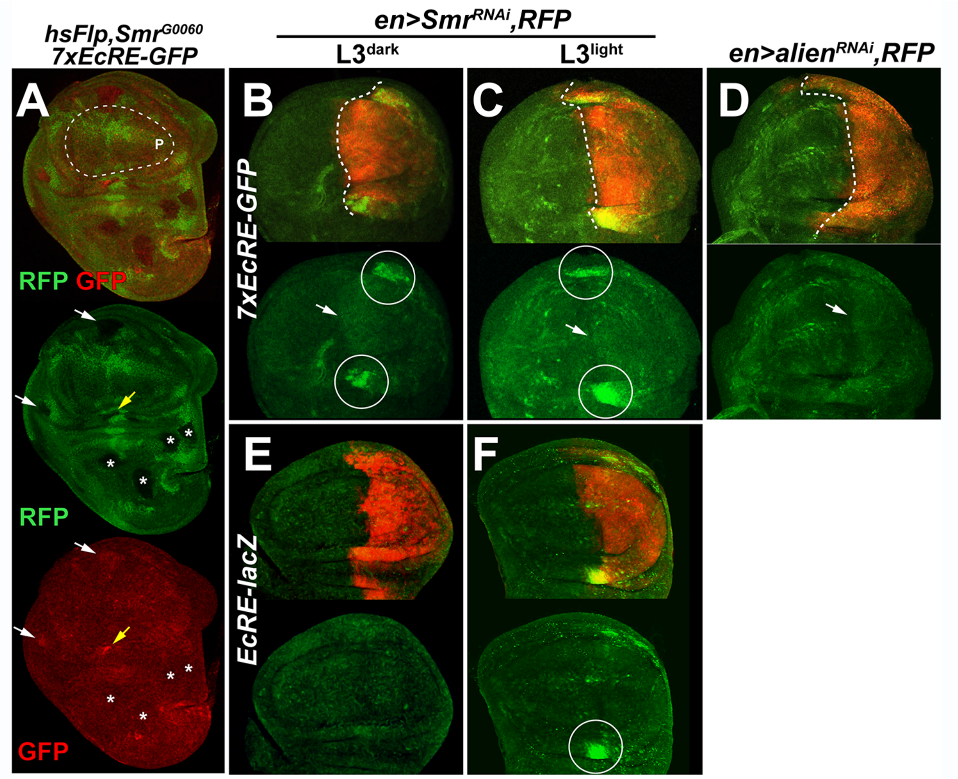


**Supplemental Figure 7. Smr$^{RNAi}$ reveals enhanced EcR-repression at the intersection of the hinge and the A:P boundary.** (**A**) Effect of *Smr*$^{G0060}$ clones (marked by absence of RFP; false colored green) on *7xEcRE-GFP* (false colored red) in an L3 wing disc. Note absence of surviving clones in the pouch (dotted line). Asterisks denote *Smr*$^{G0060}$ clones in the notum that have no effect on *7xEcRE-GFP*. Yellow arrow marks a medial hinge *Smr*$^{G0060}$ clone that activates *7xEcRE-GFP*; white arrow marks a lateral hinge clone that weakly activates *7xEcRE-GFP*. *Smr* depletion (*en>Smr*$^{RNAi}$*,RFP*) induces (**B-C**) *7xEcRE-GFP* strongly in L3$^{dark}$ and L3$^{clear}$ hinge cells at the anterior:posterior boundary (circles) and weakly across the rest of the posterior pouch cells (white arrow), (**E-F**) but only robustly induces *EcRE-lacZ* in L3$^{clear}$ cells and then only in the dorsal most part of the medial hinge (circle in **F**). (**D** )By comparison, RNAi of the EcR repressor *alien* results in mild and uniform induction of the *7xEcRE-GFP* reporter.
